# Influence of heterotrophs on phage infection of marine picocyanobacteria

**DOI:** 10.1101/2025.09.17.676900

**Authors:** Allison Coe, Sierra M. Parker, Nhi N. Vo, Sean M. Kearney, Shaul Pollak, Konnor von Emster, James I. Mullet, Kurt G. Castro, Sallie W. Chisholm

## Abstract

Picocyanobacteria *Prochlorococcus* and *Synechococcus* coexist with both their lytic phages and heterotrophic bacteria in the oceans. These lytic phages are a significant cause of mortality, and heterotrophic bacteria have been shown to increase the fitness of *Prochlorococcus* by reducing oxidative stress and cross feeding under extended darkness. Studies of *Prochlorococcus-*phage interactions are often done with xenic cultures as it has been historically difficult to obtain and maintain heterotroph-free cultures. Here we examine the effects of heterotrophic bacteria on phage infection in *Prochlorococcus* and *Synechococcus* by comparing phage infection dynamics in cultures with and without heterotrophs present. We found that *Prochlorococcus* populations resumed growth following infection only in the presence of heterotrophs, independent of phage:host or heterotroph:host ratios. In phage:host pairing with *Synechococcus* the outcomes varied, suggesting that the impact of heterotrophs on phage infection may be dependent on the phage:host interaction. In cases where the host recovered from phage infection, heterotrophs appeared to facilitate it both by mitigating oxidative stress and possibly supplying organic carbon sources, which may support post-infection growth. Furthermore, *Prochlorococcus* and *Synechococcus* populations that recovered from infection were resistant to phage infection when transferred to fresh media. Evidence argues against genetic change as the mechanism of resistance, suggesting that *Prochlorococcus* and *Synechococcus* populations in co-culture with heterotrophs undergo non-genetic adaptations during recovery from phage infection, likely driven by heterotroph-derived organic compounds that reshape host metabolism and confer protection against future lysis.

## Introduction

Marine picocyanobacteria *Prochlorococcus* and *Synechococcus* are small (~<1-2µm) primary producers in the marine community with a global population estimated at 3×10^27^ and 7×10^26^ cells, respectively (Flombaum et al. 2013). These genera are found in abundance (~10^4^ - 10^5^ cells mL^−1^) in the upper 200m of the world’s oceans and coexist with ~10^5^ - 10^6^ mL^−1^ heterotrophic bacteria and ~ 10^5^ - 10^7^ phage mL^−1^, with which they have the potential to interact (Johnson et al. 2006; Wigington et al. 2016; Follett et al. 2022; Comfort et al. 2024). *Prochlorococcus* and *Synechococcus* display high rates of photosynthetically derived fixed carbon release (Bjørrisen 1988; Johnson et al. 2006) potentially fueling up to ~40% of subtropical heterotrophic bacterial growth (Bertilsson et al. 2005). In xenic cultures, defined here as a single cyanobacterial strain coexisting with their co-isolated native heterotrophic community, without added carbon to media, the organic carbon released by cyanobacteria can support a wide diversity of heterotrophic species (Zheng et al. 2018; Kearney et al. 2021).

The presence of heterotrophs in a culture also influences the metabolism of *Prochlorococcus*. Indeed, within 5 h of introducing a heterotroph to the *Prochlorococcus* culture, the cell’s transcriptional profile shifts in ways that implicate a number of cellular activities and metabolic pathways (Biller et al. 2016). While the organic compounds produced by *Prochlorococcus* support heterotrophic growth, the heterotrophs, in turn, recycle components of the dissolved organic matter (DOM) and release metabolites that *Prochlorococcus* can utilize — creating a dynamic, mutualistic exchange cycle within the community (Biller et al. 2016, 2018; Coe et al. 2016, 2021, 2024; Wu et al. 2022). This is further supported by *Prochlorococcus’* demonstrated ability to transition to mixotrophic metabolism, facilitating heterotrophic assimilation of various compounds, including amino acids and glucose (Zubkov et al. 2004; Michelou et al. 2007; Mary et al. 2008; Muñoz-Marín et al. 2013). In addition, certain “helper” heterotrophs mitigate oxidative stress in *Prochlorococcus* — an environmental challenge that *Prochlorococcus* is particularly sensitive to as it does not produce catalase (Morris et al. 2008, 2011; Hennon et al. 2017).

*Prochlorococcus* and *Synechococcus* are vulnerable to infection by cyanophages, which outnumber cyanobacteria by 2- to 4-fold in the North Pacific, but infections are estimated to contribute only up to 5% of *Prochlorococcus’* daily mortality, suggesting successful encounters and infections are rare (Mruwat et al. 2021). On a global scale, lysed prokaryotic cells contribute ~145 Gt C per year to the DOM pool in tropical and subtropical oceans (Lara et al. 2017), representing a major flux within the carbon cycle. In the case of cyanobacteria, phage-induced lysis fuels heterotrophic growth, driving shifts in the community structure (Zheng et al. 2021; Man et al. 2024). Notably, DOM produced by viral lysis of *Prochlorococcus* is compositionally distinct and more bioavailable than what is passively released by exponentially growing *Prochlorococcus*, and its composition can vary with different phage:host combinations, leading to differences in the diversity and structure of the heterotrophic community (Xiao et al. 2021). DOM produced by *Synechococcus* viral lysates have been found to be rich in nitrogen-containing compounds (Ma et al. 2018; Zheng et al. 2021), and as the lysates are metabolized by heterotrophic bacteria, inorganic nutrients such as ammonium are remineralized (Zheng et al. 2021). As pointed out earlier, heterotrophs influence metabolic pathways within *Prochlorococcus* cells; therefore, an increase in heterotroph abundance and a change in community composition due to phage infection could further alter *Prochlorococcus* and *Synechococcus’* metabolism, potentially impacting subsequent cyanophage infection dynamics.

Numerous studies have explored pairwise phage-cyanobacteria and heterotroph-cyanobacteria interactions. Here, we take the next step in understanding community dynamics by examining their three-way dynamics by assessing whether the presence of heterotrophic bacteria in clonal cultures of cyanobacteria influences phage infection dynamics — either directly or indirectly. To this end, we compare infection dynamics in axenic (cyanobacteria only), monoxenic (cyanobacteria plus a single heterotroph, *Alteromonas macleodii* MIT1002, hereafter referred to as *Alteromonas*), and xenic (cyanobacteria plus a community of heterotrophs that were originally co-isolated with them) *Prochlorococcus* and *Synechococcus* cultures. Finding that the presence of heterotrophs promotes host survival, we next examine how the ratios of phage:host and heterotroph:host impact host recovery from phage infection. We then do a series of experiments designed to help understand how heterotrophs might infer resistance to phage infection. Finally, we investigated whether shifts in the heterotrophic community following sequential phage infections might play a role in shaping host responses, including the potential development of phage resistance.

## Materials and Methods

### Cultures

Axenic (cyanobacteria alone), monoxenic (cyanobacteria plus a single heterotroph), and xenic (cyanobacteria plus a community of heterotrophs that were originally co-isolated with them) *Prochlorococcus* and *Synechococcus* cultures were grown in 0.2 μm filtered sterile natural seawater amended with Pro99 nutrients prepared as previously described (Moore et al. 2007). Prior to addition to cyanobacteria cultures, *Alteromonas macleodii* MIT1002 cultures were grown in 0.2 μm filtered sterile natural seawater amended with ProMM nutrients as previously described (Berube et al. 2015). All cells were grown in continuous light at ~15 μmol photons m^−2^ s^−1^ in 24°C. At the beginning of each experiment, exponentially growing cultures of the cyanobacterium were enumerated by flow cytometry (described below) so that initial concentrations of *Prochlorococcus* could be set at 1 or 1.5 × 10^7^ cells mL^−1^. In cases where *Alteromonas* was added, its initial concentration was set to be 10 fold less (1 or 1.5 × 10^6^ cells mL^−1^) than that of the cyanobacterium, unless otherwise stated. This dilution was selected on the basis that, throughout exponential growth, *Alteromonas* abundances in *Prochlorococcus* co-cultures are generally an order of magnitude lower (Coe et al. 2016, 2021, 2024)(Coe et al, 2016, 2021, 2024). *Alteromonas* was chosen for these experiments because it is commonly present in *Prochlorococcus* cultures and was originally isolated from a *Prochlorococcus* culture (Biller et al. 2015; Coe et al. 2016). Heterotroph:cyanobacteria ratios in xenic cultures – i.e. those with an associated heterotrophic community – ranged from 4:1 to 10:1. The composition of these communities (except for those with *Synechococcus* MITS9451), was described in Kearney et al. (2021). Although the composition may have slightly changed since then, we note that the current communities in xenic *Prochlorococcus* MED4 cultures generally align with those previously reported.

Cells were grown in either 96 well cell culture plates (cat#353072, Corning, Glendale, AZ) or 25mm borosilicate test tubes (cat#47729-586, Avantor VWR, Radnor, PA) with 5 or 3 biological replicates, respectively. Growth was monitored via bulk chlorophyll fluorescence in plates (Synergy2, Agilent BioTek, Santa Clara, CA) or in tubes (TD700, Turner Designs, Sunnyvale, CA) and by flow cytometry (see below). For experiments involving reactive oxygen species and organic carbon additions, 5mM stocks were made with ultra-pure water and cell-culture grade chemicals (e.g., Bioxtra, Millipore Sigma, Burlington, MA). Stocks were sterilized by filtering through a 0.2µm Supor syringe filter (cat# 4506, Cytiva, Marlborough, MA) and stored at 4°C until use.

### Phages

In this study, we used three cyanophages: the podo- and T7-like specialist phage P-SSP7 and the myo- and T4-like phage P-HM2 to infect *Prochlorococcus* MED4, and the myo- and T4-like generalist phage Syn9 to infect *Synechococcus* WH8102, WH7803, and MITS9451. Phage titers were determined using the Multiple Probable Number (MPN) method as described by Jarvis et al. (2010) and phages were added at the onset of each experiment at a multiplicity of infection (MOI) of 0.1, unless otherwise noted. Phage and cyanobacteria were co-incubated for 60 min to allow for adsorption, after which Pro99 medium was added to bring the final well volume to 200 µL.

### Flow cytometry

*Prochlorococcus* cell abundances were measured as previously described (Coe et al. 2021) using a Guava 12HT flow cytometer (Luminex Corp., Austin, TX, U.S.A.) equipped with a 488 nm blue laser. Cells were analyzed based on chlorophyll autofluorescence (692/40 nm) and DNA content (530/40 nm) following staining with 1x SYBR Green I (Invitrogen, Grand Island, New York, U.S.A.) and a 60 min dark incubation prior to analysis. All flow cytometry data were processed and analyzed using FlowJo software (version 10.8.1; Flowjo LLC, BD Life Sciences, Ashland, OR, USA).

### DNA sequencing and heterotrophic community analysis

To examine genetic mutations and identify shifts in heterotroph composition across sequential infections, genomic DNA was extracted from biological triplicate cultures harvested at late exponential phase following both the first and second infection. Cells from 15 mL of culture were spun down at 7,197 *x g* for 20 min with 0.003% Pluronic F-68 Polyol (MP Biomedicals, Cat# 2750049), which enhanced cell aggregation and biomass recovery. The resulting pellets were stored at −80°C prior to DNA extraction. Genomic DNA was isolated using the protocol found in Wilson et al. (1997) modified for *Prochlorococcus* as described in Coe et al. (2025). Sequencing libraries were prepared with the Illumina NexteraXT kit (cat# FC-131-1096, Illumina, San Diego, CA) following manufacturer’s protocol. Samples were indexed using the TG Nextera® XT Index Kit v2 Set A (cat# TG-131-2001 for kit A or ending in 2 for kit B), and sequencing was performed on the Illumina NextSeq500 at the MIT BioMicro Center. Sequencing reads were deposited to SRA under BioProject PRJNA1302459 (Supporting Information Table S1).

To identify SNPs and indels before and after sequential phage infections in monoxenic and xenic *Prochlorococcus* MED4 and xenic *Synechococcus* MITS9451 cultures, we used a modified version of the variant-calling workflow described by Conwill et al. (2022). Key adjustments included setting the minimum allele frequency threshold to 0.1 and increasing the maximum read depth per position to 5000. Briefly, sequencing reads were trimmed with Cutadapt v.1.18 (Martin 2011), aligned to reference genomes *Prochlorococcus* MED4 (IMG accession Ga0065304_11) and *Synechococcus* MITS9451 (NCBI accession NZ_CP138971) using bowtie2 v2.2.6 (Langmead and Salzberg 2012), and variants were called with Samtools v1.9 and BCFtools v.1.2 (Danecek et al. 2021) (Supporting Information Table S2). Single nucleotide variants (SNVs) present in all samples were excluded from further analysis. Similarly, indels were extracted from the BCFtools output, and those shared across all treatments were also removed from downstream analysis.

To characterize the heterotrophic community co-isolated with the host cells that were serially transferred over many years, we used the ProSynTax dataset and a modified version of its associated workflow (Coe et al. 2025). Because the concentration of the added DNA standard, originally intended to obtain absolute quantification, was not recorded, we proceeded with relative quantification instead. To prevent interference from the standard, reads were processed through a two-step filtering approach, as read mapping may not detect all standard-derived reads. Reads were first trimmed using BBDuk v39.18 (Bushnell et al. 2017), then mapped to the *Thermus thermophilus ATCC BAA-163* reference genome (NCBI accession AE017221) using bowtie2 v2.5.4 (Langmead and Salzberg 2012), and standard-derived reads were removed with SeqKit v2.10.0 (Shen et al. 2024). Taxonomic classification of the resulting reads was performed with Kaiju v1.10.1 (Menzel et al. 2016) against the ProSynTax v1.1 dataset (Coe et al. 2025), which includes an extensive reference set for heterotroph taxonomy. Any remaining reads classified as *Thermus thermophilus* were filtered to ensure accurate downstream analysis. Genera that did not reach at least 1% of classified reads in either the pre- or post-infection samples were grouped under the category “Others” (Supporting Information Table S3).

## Results and Discussion

### Heterotrophic effects on Prochlorococcus phage infection

Rendering *Prochlorococcus* cultures axenic has been historically challenging due to their dependence on co-occurring heterotrophs to mitigate oxidative stress (Morris et al. 2008, 2011), and consequently, early studies of phage infection were conducted on xenic cultures (Sullivan et al. 2003; Lindell et al. 2004). With recent advances facilitating the routine isolation of axenic cultures (Berube et al. 2015; Kearney et al. 2022), we systematically investigated the role of heterotrophs in shaping infection dynamics. We grew *Prochlorococcus* MED4 under three conditions: axenic, monoxenic (referred to as “+*Alteromonas*” in figures), and xenic (cyanobacteria plus a community of heterotrophs that were originally co-isolated with them; Kearney et al. 2021). Experimental cultures were infected with podovirus P-SSP7 (Figure 1). Control cultures (“no phage control”) grew as expected, with minimal differences across the three culture types (Figure 1 a). Following phage introduction, all treatments underwent lysis, as evidenced by the flat-lining of bulk culture chlorophyll fluorescence relative to the controls for 10 days. Only cultures with heterotrophs – both xenic and +*Alteromonas* – resumed growth after ~10 days post-infection (Figure 1 b).

**Figure 1.**
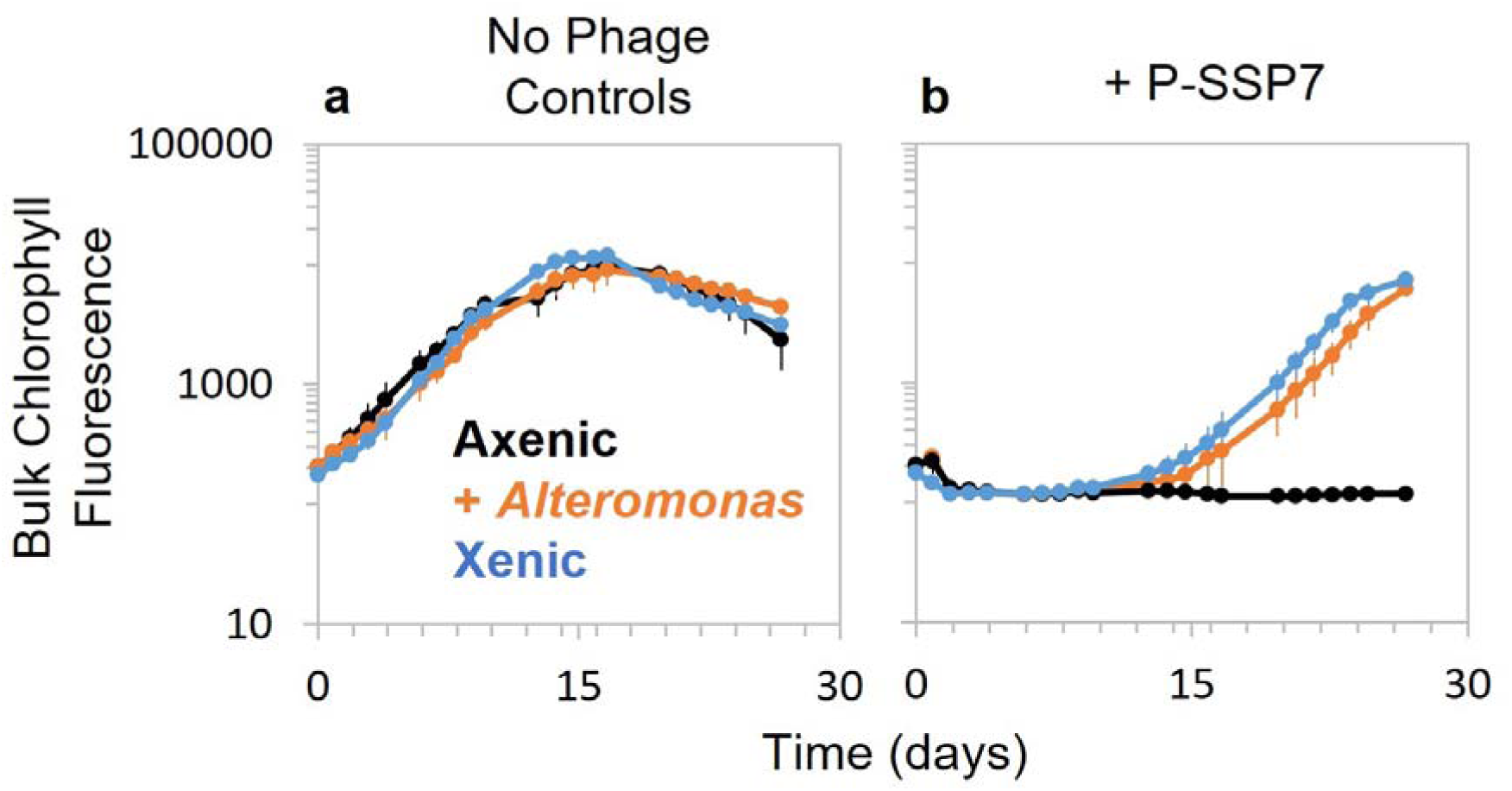
Heterotroph presence enables recovery post-phage infection of *Prochlorococcus*. Axenic (black), monoxenic (*+Alteromonas*) (orange), and xenic (community of heterotrophs) (blue) cultures of *Prochlorococcus* MED4 without (a) or with (b) cyanophage P-SSP7. Phages were added at the onset of the experiment and the culture dynamics were monitored via bulk culture chlorophyll fluorescence. Xenic cultures were maintained separately from axenic cultures, each harboring an independently maintained heterotrophic community.

This difference posed the question of whether phage had adsorbed to the heterotrophs, effectively decreasing the multiplicity of infection (MOI — i.e., ratio of phage to *Prochlorococcus* host) thus diluting phage encounters with *Prochlorococcus*. To explore this, we adjusted the multiplicity of infection — set to 0.1 in all other experiments unless otherwise specified — to see if it would influence recovery, comparing axenic and monoxenic *Prochlorococcus* MED4 cultures (Figure 2). Across a wide MOI range (10-0.001), both axenic and monoxenic cultures showed similar infection dynamics (+/− 2 days), while MOIs below 0.0001 failed to establish infection in either culture (Figure 2). Only the cultures containing *Alteromonas* recovered post-infection and the timing of recovery was consistent across all effective MOIs (Figure 2 b). Notably, in the low-MOI cultures that failed to lyse, *Alteromonas* also prolonged *Prochlorococcus* survival into the post-stationary (decline) phase compared to axenic cultures, which rapidly collapsed, consistent with observations by Weissberg et al. (2022) who suggested that these two strains exhibit mutualistic interactions. Thus, recovery only occurs in the presence of heterotrophs and is independent of the initial phage-to-host ratio, suggesting that it is driven by heterotroph-associated processes rather than by the magnitude of the initial infection.

**Figure 2.**
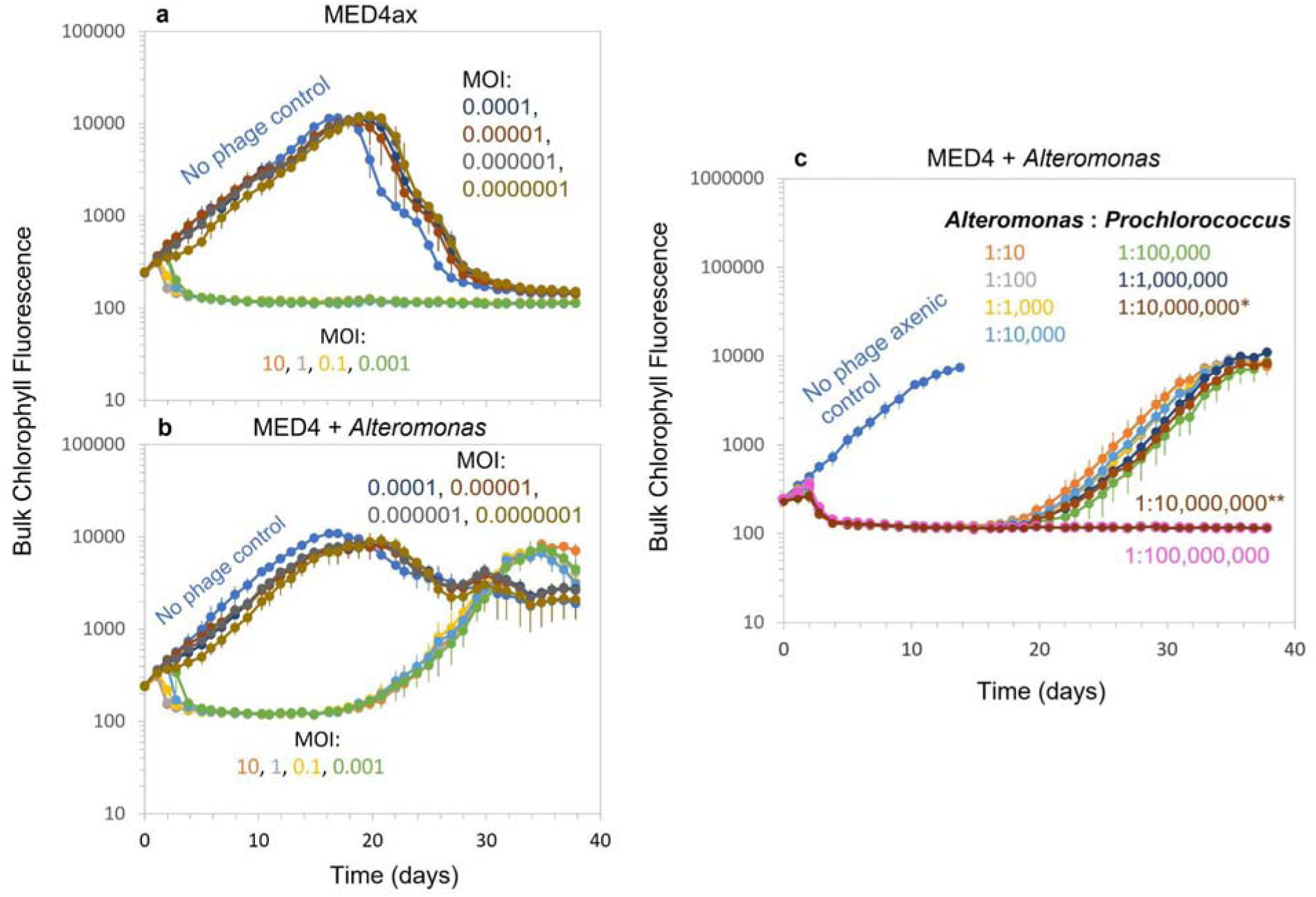
Phage:host ratio (MOI) and heterotroph:host ratio at onset of experiment does not alter recovery of host from phage infection. (a) Axenic and (b) monoxenic (*+Alteromonas*) *Prochlorococcus* MED4 cultures infected with cyanophage P-SSP7 at the onset of the experiment using MOIs (phage:host ratio) ranging from 10 to 0.0000001 and (c) *Prochlorococcus* MED4 grown with and without *Alteromonas* at *Alteromonas*:*Prochlorococcus* ratios from 1:10 - 1:10,000,000. In the 1:10,000,000 ratio (brown), one out of the 6 biological replicates resumed growth (*) while the remainder did not resume growth (**). The 1:100,000,000 ratio (pink), effectively zero, served as a negative control for *Alteromonas* presence. All cultures - except the no phage control (blue) - were infected with P-SSP7 at the onset of the experiment and the infection dynamics were monitored via chlorophyll fluorescence.

As the timing of recovery is decoupled from the host:phage ratio, we wondered whether the concentration of heterotrophs at the onset of the experiment would have an influence on these dynamics. What is the minimum concentration of heterotrophs to facilitate recovery after infection? To test this, we grew axenic *Prochlorococcus* MED4 with varying initial ratios of *Alteromonas*:*Prochlorococcus* cells, ranging from 1:10 - 1:100,000,000. The actual starting *Prochlorococcus* abundance was 1.5 × 10^7^, while *Alteromonas* concentrations ranged from 1.5 × 10^6^ to 1.5 × 10^−1^ cells mL^−1^, with the higher concentration representing the standard 10-fold dilution used in all experiments and the lowest serving as the control for absence of *Alteromonas*. Cultures were then infected with P-SSP7 (Figure 2 c). *Prochlorococcus* resumed growth at similar times across all treatments where heterotrophs were present, regardless of the initial *Alteromonas*:*Prochlorococcus* ratio (Figure 2 c, days 17-18). The only exception was the 1:10,000,000 ratio, where only 1 out of the 6 replicates resumed growth (Figure 2 c brown lines). This anomaly could be attributed to the inherent errors that come with pipetting samples of cells with such low abundance. Overall, these results show that as long as heterotrophs are present – even at one cell per 10,000,000 *Prochlorococcus – Prochlorococcus* can recover post-infection. Thus, the heterotrophs are either important for phage resistance or recovery post-infection, irrespective of MOI or heterotroph abundance.

### Heterotroph abundance during phage infection

When heterotrophs are introduced to *Prochlorococcus* cultures maintained in natural seawater-based media without added organic compounds, they can grow initially, fueled by background seawater organic matter. Once this is depleted, however, heterotroph growth is entirely dependent on photosynthate released by *Prochlorococcus*, limiting their proliferation (Coe et al. 2016, 2024; Becker et al. 2019). Evidence also shows that phage-induced lysis of *Prochlorococcus* and *Synechococcus* can supply a sudden influx of organic matter, supporting increased heterotrophic growth (Xiao et al. 2021; Man et al. 2024). While bulk chlorophyll fluorescence data in Figures 1 and 2 reflect cyanobacteria dynamics, they do not capture heterotrophic growth, leaving it unclear whether organic matter from lysed *Prochlorococcus* cells influences heterotrophic growth in these co-cultures. Given the central role of heterotrophs on nutrient recycling and microbial community structure (Aharonovich and Sher 2016; Biller et al. 2018; Follett et al. 2022; Wang et al. 2022), changes in their abundance during infection raise the possibility of indirect effects, through cross-feeding, altered MOI, or other community-level feedbacks, that could shape phage:host interactions. To examine this, we used flow cytometry to measure abundance of both populations throughout infection and subsequent recovery in xenic *Prochlorococcus* MED4 cultures (i.e., cultures containing a heterotrophic community) infected with phage P-SSP7, allowing us to examine host-heterotrophs dynamics under conditions more representative of natural communities. *Prochlorococcus* abundance initially rose for one day, then declined due to lysis until day 5, when growth resumed (Figure 3, blue line). During this lysis phase, heterotroph abundance surged - doubling more than once per day over the first 5 days (Figure 3, orange line). As *Prochlorococcus* began to recover, heterotroph growth slowed markedly, nearly stalling (Figure 3, orange line). Once *Prochlorococcus* abundance surpassed that of the heterotrophs, heterotroph growth resumed again (Figure 3, orange line).

**Figure 3.**
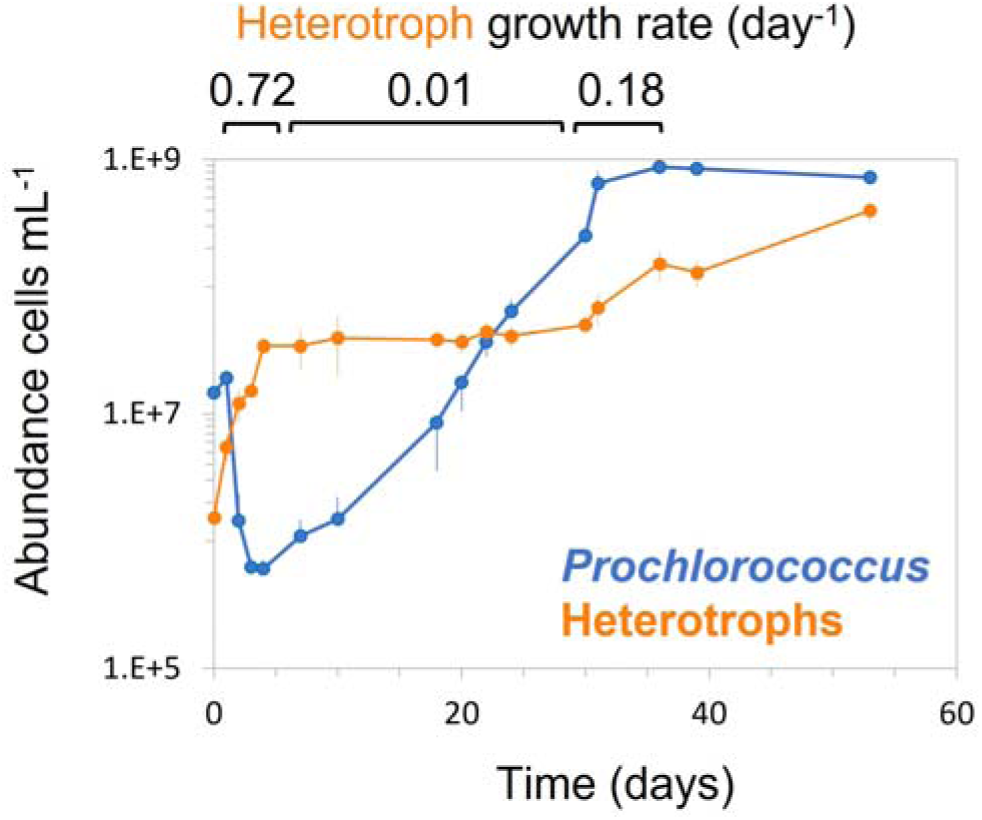
Phage infection causes *Prochlorococcus* decline and a transient surge in heterotroph abundance. Abundance of xenic *Prochlorococcus* MED4 (blue) and their associated community of heterotrophs (orange) after infection with podo-like virus P-SSP7 at the onset of the experiment. Heterotroph growth rate (day^−1^) during lysis and after *Prochlorococcus’* resumption of growth is indicated in black above the figure.

To confirm the initial spike in heterotrophic growth was due to lysis, we conducted a parallel experiment comparing heterotroph abundance in xenic *Prochlorococcus* MED4 cultures with and without P-SSP7 (Supporting Information Figure S1). In the no phage control, heterotroph abundance increased after the first day, plateaued until day 4, and then increased again once *Prochlorococcus* abundance surpassed that of the heterotrophs (Supporting Information Figure S1, black lines). In contrast, phage-infected cultures showed a significant increase in heterotroph abundance during the first three days (*t-test* p = < 0.05 or 0.01), before leveling off at day 4, just as *Prochlorococcus* recovery began (Supporting Information Figure S1 b, blue lines). These findings suggest that heterotrophs rapidly respond to organic matter released during lysis, and this transient surge may play a role in shaping phage infection and recovery dynamics.

### Interaction timing between Prochlorococcus and heterotrophs

Timing of recovery is not a function of phage:host ratio or initial heterotroph:host ratio. However, we observed a significant increase in heterotroph abundance during infection (Figure 3, Supporting Information Figure S1), prompting us to wonder whether this early heterotroph increase, potentially influencing host recovery through mechanisms such as cross-feeding, plays a role. Specifically, we investigated if the timing of heterotroph introduction, including the effects of early exposure or “pre-conditioning”, would reveal how long heterotrophs must be present to affect phage lysis and recovery dynamics. Unlike previous experiments that added heterotrophs only at infection onset, we varied the timing of heterotroph addition to determine if recovery depends on early presence or if later introduction can still support it. To test this, we compared axenic and monoxenic *Prochlorococcus* MED4 cultures infected with phage P-SSP7, in which *Alteromonas* was added at specific time points: the onset of the experiment (T0), after 2 days (T2), or after 5 days (T5) (Figure 4, black, blue, and red ticks on the axis, respectively). As a control, fresh Pro99, the same media the cells were grown in and the heterotrophs diluted into, was added to the axenic and xenic controls.

**Figure 4.**
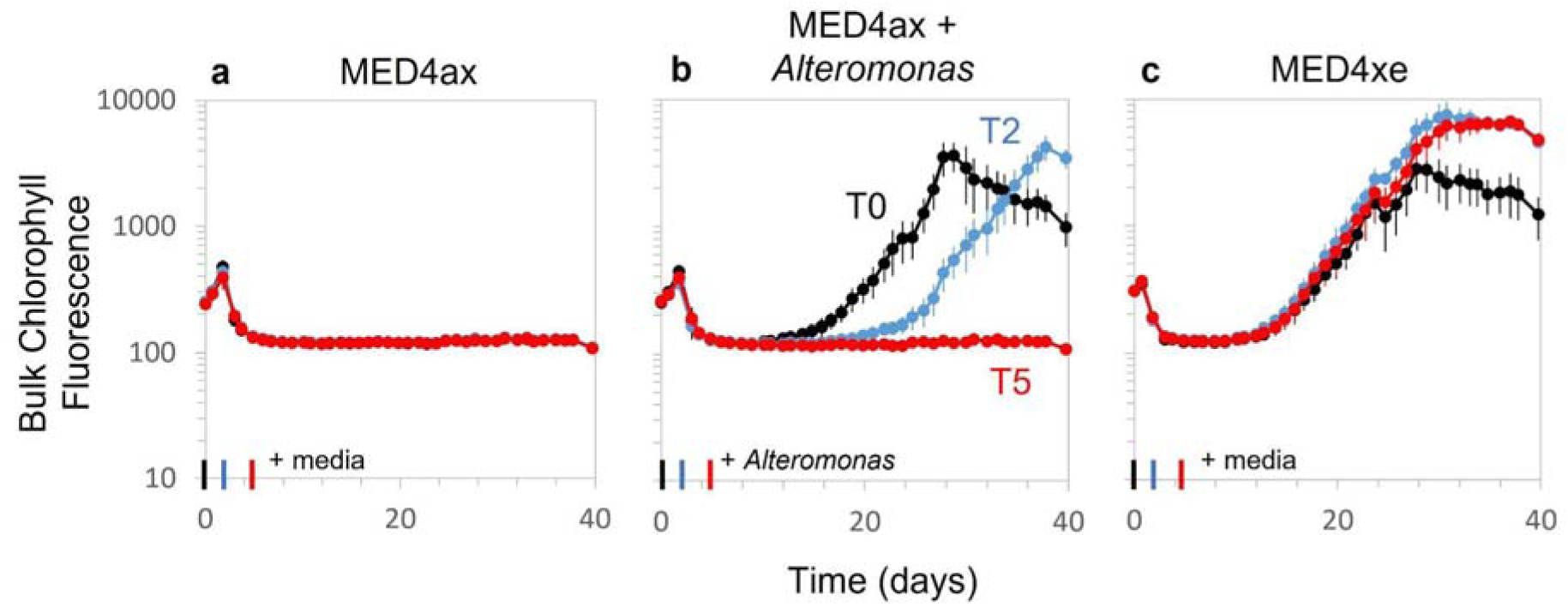
Timing of *Alteromonas* addition influences phage infection dynamics. (a) Axenic (“ax”), (b) monoxenic (+*Alteromonas*), and (c) xenic (community of heterotrophs) *Prochlorococcus* MED4 infected with podo-like phage P-SSP7 at the onset of the experiment. Additions of equivalent volume of media or *Alteromonas* (1 × 10^6^ cells mL^−1^) were added (as indicated by tick marks on x-axis) to cultures at T0 (black), T2 (blue), and T5 (red) and the infection dynamics were monitored via bulk chlorophyll fluorescence.

The timing of heterotroph addition, creating monoxenic cultures, had a significant influence on recovery time (Figure 4 b). When *Alteromonas* was added at T0, recovery proceeded as expected (Figure 4 b, black line). Addition at T2 also supported recovery, though with a noticeable delay (Figure 4 b, blue line). However, when *Alteromonas* was introduced at T5, *Prochlorococcus* failed to resume growth (Figure 4 b, red line). This suggests that in monoxenic cultures a critical window for heterotroph-mediated population recovery exists between days 2 and 5 post-infection — possibly before the cyanobacterial population is fully lysed — and that *Prochlorococcus* must be in co-culture with heterotrophs for a minimum duration within this period to develop effective resistance or support for recovery.

To further investigate the effect of prior exposure to heterotrophs (i.e., “pre-conditioning”), we compared these outcomes with xenic cultures, which contained a natural, well-established community of heterotrophs (Figure 4 c). In xenic cultures, *Prochlorococcus* consistently resumed growth following phage infection, with recovery dynamics comparable to the T0 monoxenic treatment (Figure 4 b and c). However, the media additions, included to control for the same nutrient inputs received by the monoxenic cultures, extended the duration of exponential growth after infection, likely due to the nutrient enrichment rather than biotic interaction (Figure 4 c, blue and red lines). These results reinforce the idea that the timely presence of heterotrophs - whether already well-established in the community through preconditioning, introduced at the onset of infection, or added within a critical window - can facilitate host survival and population rebound following phage infection. Therefore, heterotrophs provide critical support to *Prochlorococcus* during phage infection, but the mechanism is unclear. To investigate this further, we examined how heterotrophs influence infection dynamics across different phage:host combinations.

### Heterotrophic influence across different host:phage systems

To begin to explore whether heterotroph-mediated recovery from phage infection is a general feature of cyanobacteria-phage interactions, we examined additional phage-host combinations. Specifically, we cultured *Prochlorococcus* MED4 and *Synechococcus* WH8102 under axenic, monoxenic, and xenic conditions. In addition, *Synechococcus* WH7803 was tested in both axenic and xenic cultures, and *Synechococcus* MITS9451 was tested in xenic cultures. Each culture was infected with either podovirus- or myovirus-like phages (P-SSP7, P-HM2, or Syn9) at the onset of the experiment. MED4 + P-SSP7 data from Figure 1 were included for direct comparison. Control cultures without phage infection grew similarly across treatments (Figure 5 a, c, e, g, i). Upon phage infection, all cultures exhibited lysis within 4 days, and all cultures containing heterotrophs – except *Synechococcus* WH8102 + *Alteromonas* and xenic WH7803 – resumed growth (Figure 5), suggesting that heterotrophs often, but not always, facilitate recovery.

**Figure 5.**
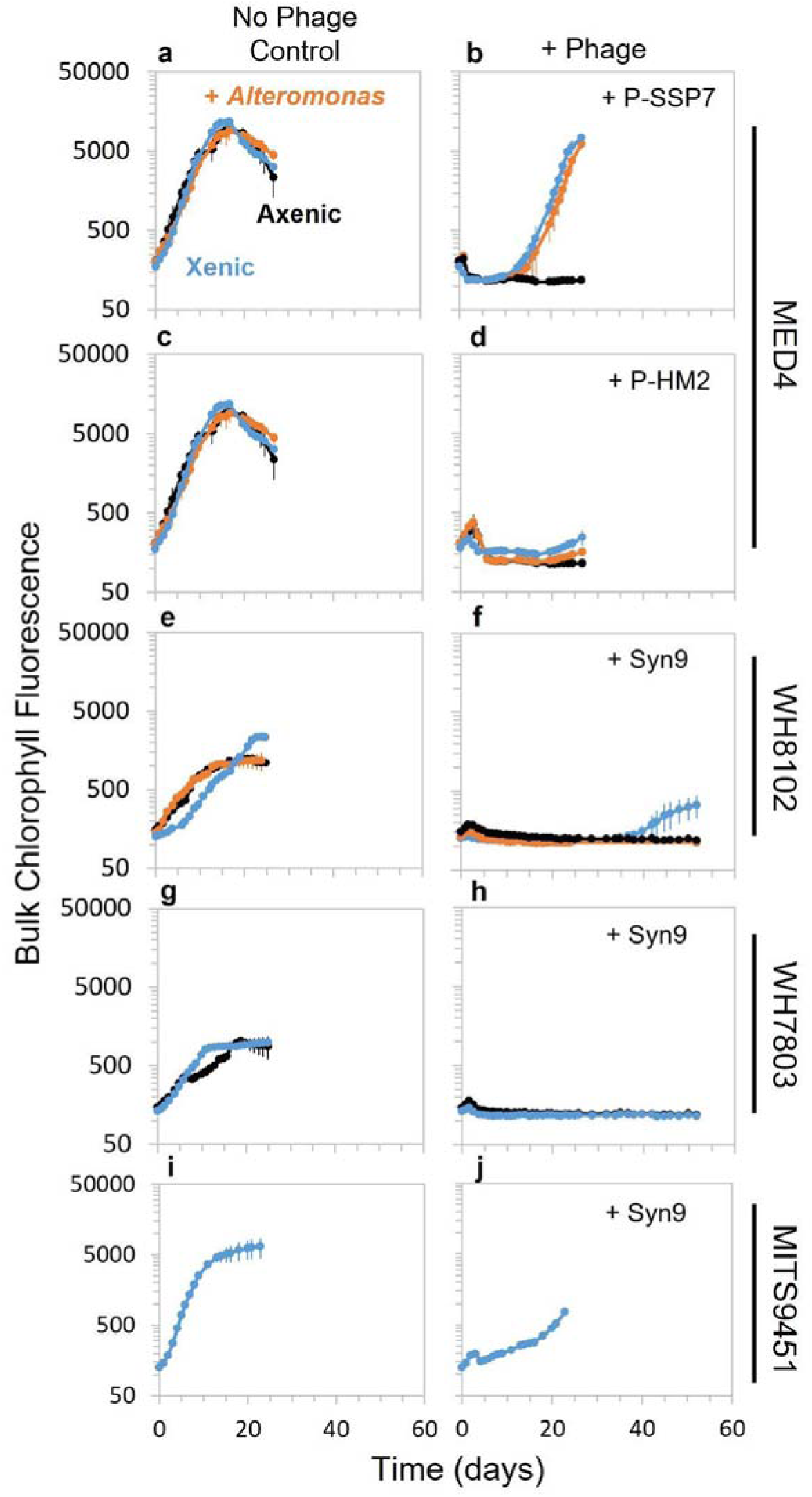
Heterotrophs facilitate recovery in multiple host-phage systems, with exceptions. Axenic (black), monoxenic (*+Alteromonas*) (orange), and xenic (community of heterotrophs) (blue) of *Prochlorococcus* MED4 without (a, c) or with (b, d) cyanophage P-SSP7 or P-HM2 or *Synechococcus* WH8102, WH7803, and MITS9451 without (e, g, i) or with (f, h, j) cyanophage Syn-9. Phages and *Alteromonas* were added at the on-set of the experiment and the infection dynamics were monitored via chlorophyll fluorescence. Xenic cultures were maintained separately from axenic cultures, each harboring an independently maintained heterotrophic community.

Infection outcomes varied across host-phage combinations. Regrowth of *Prochlorococcus* MED4 infected with P-HM2 was markedly slower than with P-SSP7 (Figure 5 b, d) and *Prochlorococcus* MED4 (with P-HM2) and *Synechococcus* WH8102 (with Syn9) exhibited reduced carrying capacities (Figure 5 d and f). Both P-HM2 and Syn9 are myovirus-like, with longer contractile tails, whereas P-SSP7 is podovirus-like, with a short tail. These morphological differences along with their larger genomes (Sullivan et al 2005, 2010), may contribute to the observed variation in host recovery. Although P-HM2 produces fewer extracellular phage per infected cell, has a shorter lytic cycle than P-SSP7 (Laurenceau et al 2020), which together with potential genome-encoded effects on host physiology, may result in slower host regrowth and reduced carrying capacity. Despite these differences, most cultures containing heterotrophs ultimately resumed growth.

Given the lack of recovery observed in monoxenic *Synechococcus* cultures in Figure 5, we wondered whether this could be due to the initial *Synechococcus*:*Alteromonas* ratio or the absence of preconditioning (i.e., prior co-culture resembling more xenic-like conditions). To test this, we compared axenic and monoxenic *Synechococcus* WH8102 cultures with treatments that varied the starting *Synechococcus*:*Alteromonas* ratio or included pre-conditioning with *Alteromonas* prior to the experiment. Cultures were grown with and without infection by Syn9 (Supporting Information Figure S2). Consistent with earlier results (Figure 5), none of the infected cultures resumed growth under any condition, indicated that neither the starting ratio of *Synechococcus* to *Alteromonas* nor preconditioning with heterotrophs was sufficient to support recovery after phage infection in *Synechococcus*. These results underscore that phage infection dynamics and recovery trajectories are strain- and phage-specific, highlighting that recovery post-infection is shaped by the specific microbial partners involved.

### Genetic variation in co-cultures

Genetic mutations conferring resistance to phage are a well-documented phenomenon in microbial systems (McGee et al. 2023; Mi et al. 2023), including in cyanobacteria (Stoddard et al. 2007; Avrani et al. 2011). To explore the mechanisms driving *Prochlorococcus* and *Synechococcus* recovery when heterotrophs are present, we first investigated whether this recovery was the result of selection for phage-resistant mutants within the host population after an infection event. To test this, we grew monoxenic and xenic *Prochlorococcus* MED4 and xenic *Synechococcus* MITS9451 with and with cyanophages (P-SSP7 and Syn9, respectively) over two consecutive infection cycles, transferring recovered cells to fresh media after the first infection to assess whether resistant variants emerged (Figure 6). All three treatments underwent a lysis phase 2-3 days into the first infection, followed by recovery (Figure 6, orange lines). During the second infection, only the monoxenic *Prochlorococcus* culture exhibited visible lysis or growth inhibition, likely due to cells transitioning into stationary phase before the transfer, while the other treatments showed no signs of reinfection (Figure 6, blue lines). Moreover, an additional phage dose added during the recovery phase of the second infection in *Synechococcus* cultures did not induce lysis (Figure 6 c, blue line). This pattern suggests that resistance to phage infection developed and persisted in the population after recovery, potentially reflecting the emergence of resistant phenotypes.

**Figure 6.**
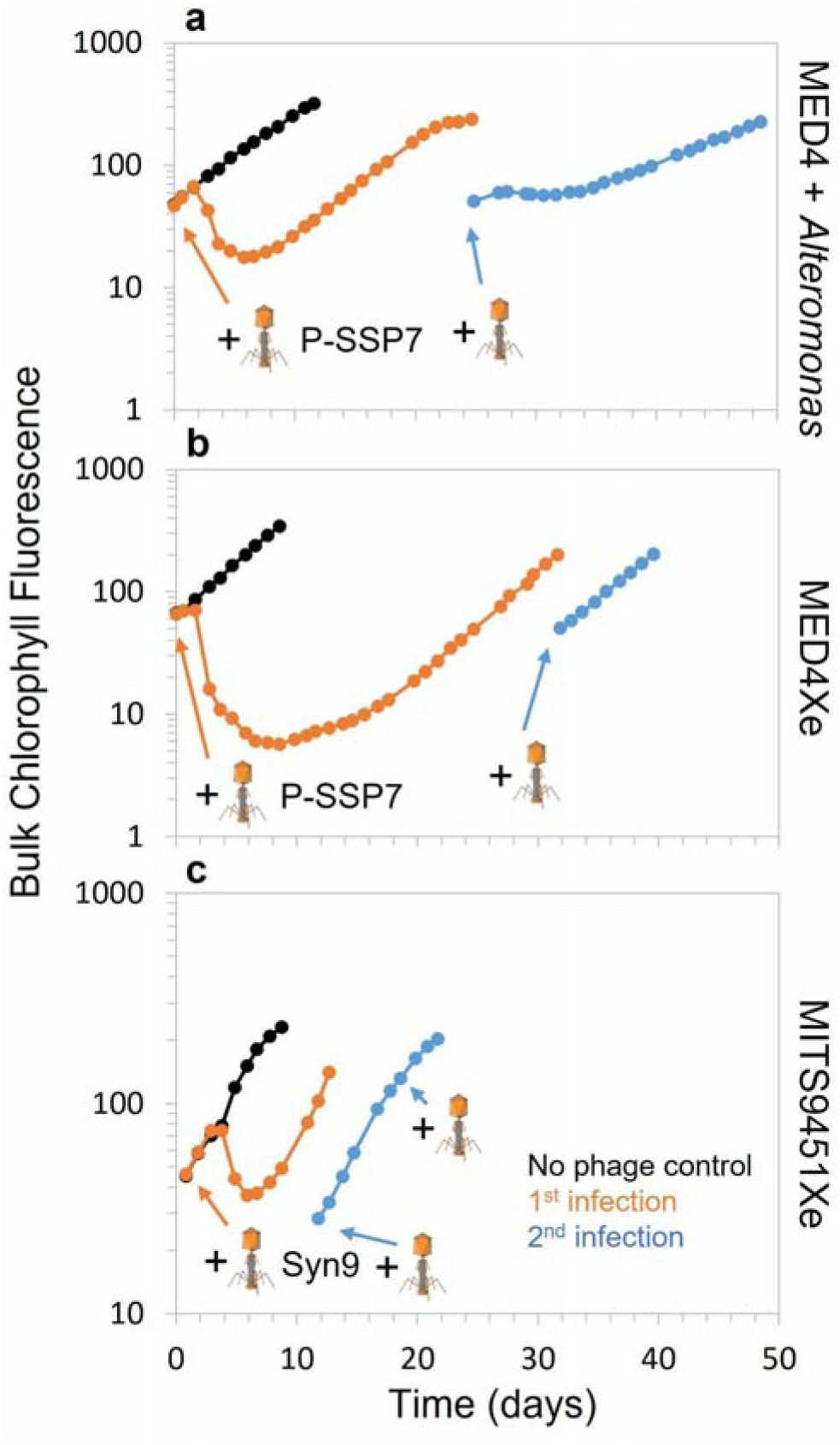
Monoxenic and xenic cultures recover from and resist consecutive phage infections. Growth of monoxenic (a) and xenic (b) *Prochlorococcus* MED4 and xenic (c) *Synechococcus* MITS9451 without (no phage control, black lines) and with cyanophages (P-SSP7 and Syn9, orange and blue lines) after consecutive infections and subsequent recovery. Cultures were transferred between the first and second infection to prevent nutrient limitation. Phages were added at the onset of each experiment, with a second addition added to *Synechococcus* MITS9451 in late exponential growth phase during the second infection (blue line).

To investigate potential mutations driving resistance, we sequenced all cultures after recovery and compared them to the no phage controls. Mutations were considered relevant if they appeared in at least two of the three biological replicates and were either unique to controls or to post infection samples. Although we cannot completely rule out the possibility that independent mutations may contribute to similar phenotypes, no single mutation was shared across all three independent cultures. Notably, there was no evidence of genetic differences including SNPs and indels, in the xenic *Prochlorococcus* MED4 and *Synechococcus* WH8102 post-infected cells (Supporting Information Table S2). While there were also no SNPs found in monoxenic *Prochlorococcus* MED4, an in-frame 3 base pair insertion resulting in the addition of a valine residue at position 100 on the protein encoded by the *prs* (ribose-phosphate pyrophosphokinase) gene — a gene involved in nucleotide biosynthesis — was observed in all but one biological replicate during the first infection, and was present in all replicates following the second infection (Figure 7 and Supporting Information Table S2). While the *prs* gene is not a known phage resistance gene, its mutation may subtly modulate host metabolism in a way that influences phage replication efficiency. The rarity of this mutation across all tested host-phage combinations, and its exclusive occurrence in the presence of a single heterotrophic partner, *Alteromonas*, suggests the possibility that it arose from host-heterotroph metabolic interactions rather than as a direct consequence of phage infection. Supporting this, *prs* expression has been shown to increase in *Prochlorococcus* MED4 when co-cultured with a similar *Alteromonas* strain, relative to when it was grown alone, suggesting that the *prs* gene is somehow involved in interactions with this heterotroph (Aharonovich and Sher 2016). Further supporting this indirect effect, no mutations were found in either the xenic cultures, and no lysis occurred after reintroducing phage during mid-to-late exponential growth in MITS9451 (Figure 6). This suggests that resistance in these cultures is largely phenotypic. We propose that the broader microbial community, through increased heterotroph abundance and interactions such as metabolite exchange or signaling, alters host metabolism in ways that allow some cyanobacteria to survive phage infection. This community-mediated resistance likely reflects phenotypic plasticity, where dynamic, reversible physiological changes — rather than genetic changes — enable host survival and prevent population collapse.

**Figure 7.**
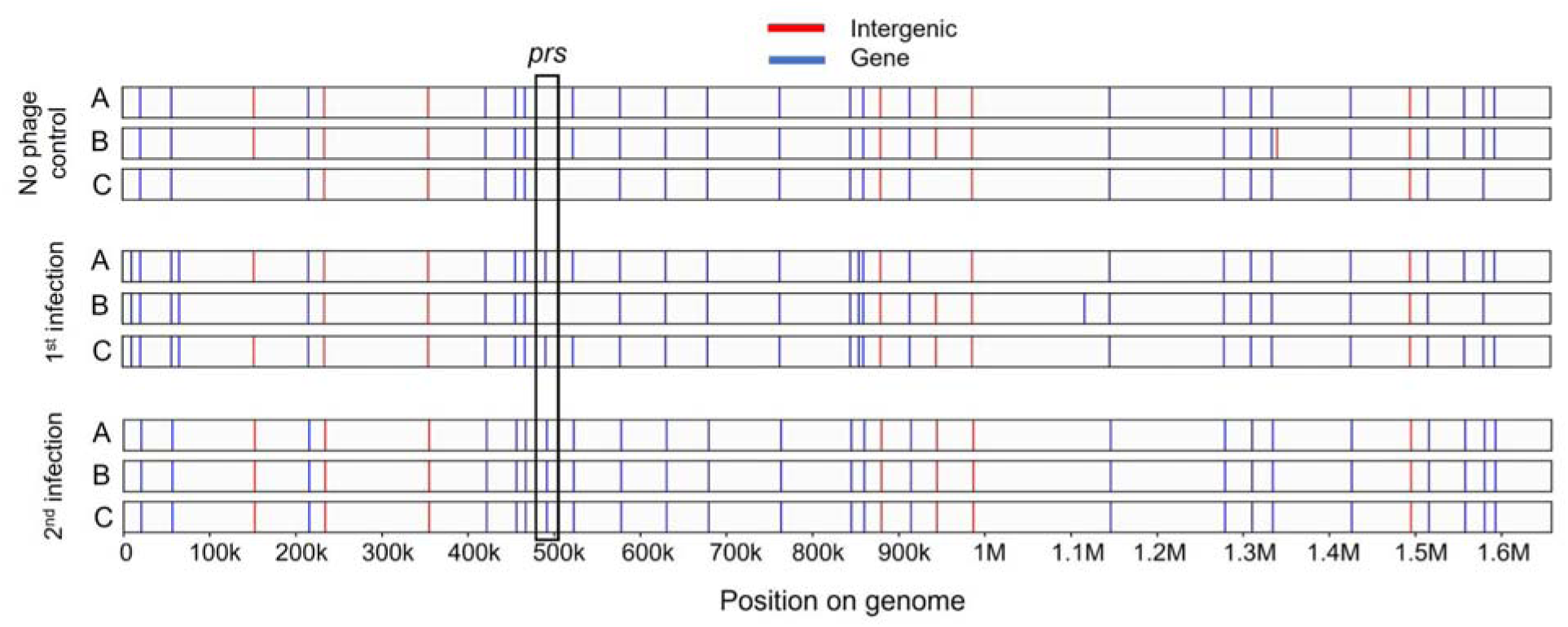
Single mutation found after consecutive phage infections in monoxenic *Prochlorococcus*. SNPs and indels located within genes (blue) or intergenic regions (red) are indicated across the genomes of *Prochlorococcus* MED4 under three conditions: before infection (no phage control), after recovery from the first infection, and after recovery from a second infection. To be deemed significant, a mutation had to be found at least 2 of the 3 replicates relative to the control, and following both the 1st and 2nd infection. The only mutation fitting these criteria was an in-frame 3 base pair insertion in the *prs* gene, highlighted with a black box.

### Non-genetic mechanisms of phage resistance

We considered several non-genetic hypotheses for how host populations recover from phage infection in the presence of heterotrophs. The first is non-specific adsorption of phages to non-hosts (e.g. heterotrophs), which would lower the effective infection rate (Murray and Jackson 1992; Weinbauer 2004). However, under the phage:host combinations tested here, we did not observe consistent recovery in *Synechococcus* when co-cultured with one or more heterotrophs at varying host:heterotroph ratios (Figure 5 and Supporting Information Figure S2). A second hypothesis concerns differences in phage infection dynamics, such as differences in adsorption kinetics, length of latent period, burst size, and extracellular decay rate, which can influence the efficiency and timing of subsequent infections (Maidanik et al. 2022). Yet this explanation seems unlikely, as *Synechococcus* WH8102 showed recovery in the presence of a heterotrophic community but not with a single heterotroph (Figure 5), even though the only experimental difference was the identity of the heterotrophs.

Having ruled out these two explanations, we next consider hypotheses focused on heterotroph-mediated effects on host physiology. One is that “helper” heterotrophs alleviate oxidative stress by leaking catalase into the surrounding medium, thereby reducing extracellular reactive oxygen species (ROS), a function *Prochlorococcus* cannot perform as it lacks the catalase gene, *katG* (Morris et al. 2008). This could promote *Prochlorococcus* recovery and potentially lower susceptibility to subsequent phage infections. Another hypothesis is that heterotroph-derived metabolites could provide nutritional support during stress (Coe et al. 2016, 2021, 2024; Biller et al. 2018; Wu et al. 2022). To distinguish between these hypotheses, we tested whether *Prochlorococcus* could recover without heterotrophs by supplementing cultures with ROS scavengers, an approach particularly relevant when *Prochlorococcus’* abundance is low and cells are more susceptible to oxidative stress (Morris et al. 2008, 2011). We also considered prior findings that, under different stress conditions (e.g. prolonged darkness), *Prochlorococcus* survival is enhanced by both ROS scavengers and organic carbon supplementation (Coe et al. 2016). We therefore asked whether recovery after infection might similarly depend on heterotroph-mediated oxidative stress relief, or alternatively, whether organic carbon released by the increased abundance of heterotrophs following lysis could alter host metabolism in ways that reduce susceptibility to phage. For example, studies with *Staphylococcus* and *E.coli* cells suggest that growth on different carbon sources can modulate metabolism, resulting in increased susceptibility, resistance to phage infection, or receptor expression changes (Weissbach and Jacob 1962; Kato et al. 1970; Schwartz 1976; Vadym and Volodymyr 2025; Marantos et al. 2025).

To examine this, axenic *Prochlorococcus* MED4 cultures were treated with either inorganic (thiosulfate) or organic (pyruvate) ROS scavengers, since we aimed to test both a non-carbon and carbon-based ROS scavenger. Additional treatments included pre-conditioned pyruvate (added to culture in the previous transfer and at the onset of the experiment), a range of carbon sources (e.g., glucose, acetate, lactate, glycerol), and a combination of pyruvate and glucose. These were compared to axenic cultures and axenic culture supplemented *Alteromonas*, both with and without cyanophage P-SSP7. As expected, the axenic cultures failed to resume growth post lysis (Figure 8 e). All ROS scavenger treatments supported host recovery following phage infection, with pre-conditioning with pyruvate delaying the onset of lysis and achieving a higher carrying capacity compared to other treatments (Figure 8 f). Among the tested organic carbon sources, only lactate enabled recovery, while acetate, glycerol, and glucose alone did not (Figure 8 g). Cultures supplemented with a combination of glucose and pyruvate had post-infection dynamics nearly identical to the axenic + *Alteromonas* treatment (Figure 8 h). Thus, pyruvate and specific carbon supplementation, particularly lactate or a combination of pyruvate and glucose, enhances *Prochlorococcus* recovery from infection with P-SSP7 by mitigating ROS and supporting host metabolism.

**Figure 8.**
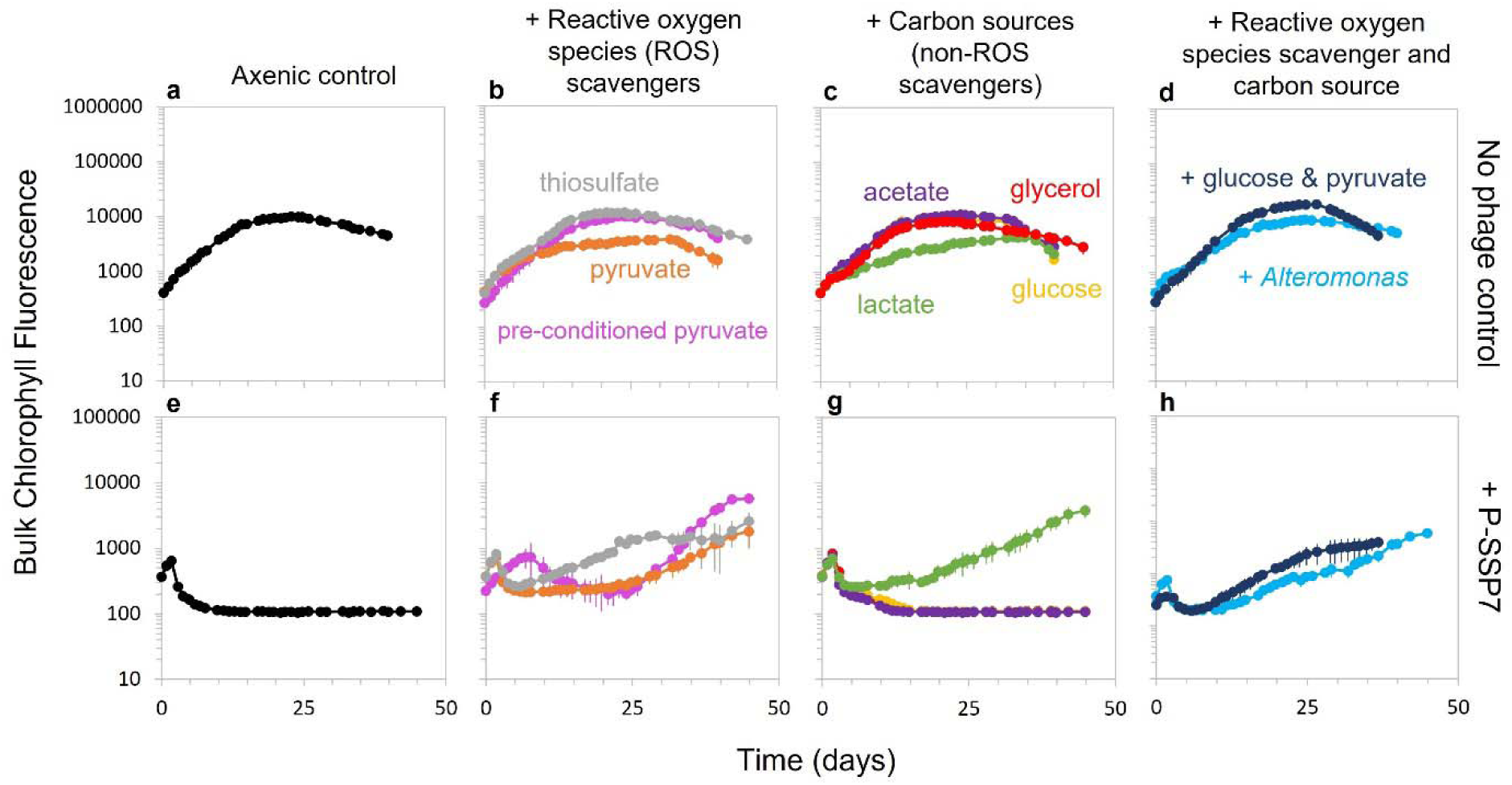
Reactive oxygen species (ROS) mitigation and carbon supplementation enable post-infection recovery. Cultures of *Prochlorococcus* MED4 were grown alone (a and e; black lines), with ROS substrates (b and f; purple, gray, and orange lines), with carbon sources (c and g; green, yellow, medium blue, red lines), or with substrates or heterotrophic bacteria that can mitigate oxidative stress and provide carbon substrates (d and h; aqua and navy blue lines). These no phage control cultures were grown in parallel with cultures infected with phage P-SSP7 at the onset of the experiment. All substrates or bacteria were added at the onset of the experiment, with the exception of the cultures that were pre-conditioned with pyruvate during a previous transfer (b, f; purple lines).

To assess whether this trend held for a different phage:host combinations, we repeated the experiment comparing axenic and monoxenic *Prochlorococcus* MED4 infected with P-HM2 and axenic and xenic *Synechococcus* WH8102 and WH7803 infected with Syn9 and compared these to axenic cultures supplemented with ROS scavenger pyruvate, organic carbon glucose, or a combination of the two. All axenic cultures infected with phage failed to recover (Supporting Information Figure S3 e, m, u). In *Prochlorococcus* MED4, pyruvate alone treatment showed brief recovery signals that could not be sustained and glucose alone resulted in a stalled population decline for ~10 days (Supporting Information Figure S3 f and g), however, the combination of these treatments enabled regrowth nearly identical to the +*Alteromonas* treatment (Supporting Information Figure S3 h). In contrast, as shown in Figure 5 and Supporting Information Figure S2, *Synechococcus* WH8102 — which also lacks *katG* and may therefore be more sensitive to oxidative stress — did not recover when supplemented with pyruvate, glucose, or both (Supporting Information Figure S3 n-p). WH8102 failed to resume growth when co-cultured with *Alteromonas* alone (Figure 5 f), but was able to resume growth with a community of heterotrophs (Supporting Information Figure S2 p), suggesting that a broader suite of metabolites, supplied by diverse heterotrophs, may be necessary to alter its metabolism in a way that allows the population to resist phage infection. WH7803 did not recover when provided with any ROS scavenger or carbon sources, even when these were supplied via heterotroph-derived supplementation, suggesting that its metabolic constraints, possibly a more limited metabolic adaptability, prevent post-infection recovery (Supporting Information S3 x).

These findings highlight that recovery dynamics differ across phage:host systems and may depend on host-specific physiological traits or interactions among microbes in the community. In *Prochlorococcu*s, ROS mitigation, particularly through pre-conditioning, and the availability of key metabolites can substitute for heterotroph-mediated recovery. The consistent success of the glucose + pyruvate combination across both phage types in *Prochlorococcus* suggests that *Alteromonas* (or other heterotrophs) may support host recovery by providing both oxidative stress relief and carbon supplementation, consistent with our findings that *Prochlorococcus* requires heterotroph-derived metabolites to survive extended darkness (Coe et al. 2016, 2021, 2024; Biller et al. 2018). While *Synechococcus* WH8102 or WH7803 did not recover under similar carbon supplementation, WH8102 did recover (as well as xenic MITS9451) in the presence of a heterotrophic community (Figure 5), indicating that such interactions can support recovery in some, but not all, *Synechococcus*:phage combinations. Although we cannot completely rule out genetic differences between axenic and xenic *Synechococcus* WH8102 and WH7803 strains, which are maintained separately and were not sequenced, prior evidence discussed above suggests this is unlikely to explain the contrasting recovery outcomes. The recovery patterns of *Prochlorococcus* and *Synechococcus* argue against a single, universal mechanism, whether phage-mediated or simply the mitigation of oxidative stress by heterotrophs, and instead suggest that recovery depends on host-specific physiology, cross-feeding interactions with specific members of the heterotrophic community, and the host’s metabolic adaptability.

### Heterotroph composition after multiple phage infections

Infected but intact *Synechococcus* cells release compounds capable of attracting heterotrophic bacteria (Henshaw et al. 2024), suggesting that phage infection may actively restructure host-associated communities by influencing which heterotrophs are recruited to the vicinity of infected cells. Given that certain metabolites appear to support recovery of some host populations after phage infection, we next investigated whether the composition of the heterotrophic community in xenic cultures shifts across successive rounds of phage infection — potentially altering the available metabolites for the host. We used Illumina shotgun metagenomic sequencing data (Figure 6) to analyze the heterotrophic communities in the xenic *Prochlorococcus* MED4 and *Synechococcus* MITS9451 cultures using ProSynTax and associated workflow for classification (Coe et al. 2025) (Supporting Information Table S3). We first examined taxonomic shifts at the class and genus level in xenic *Prochlorococcus* MED4 cultures and found significant changes in the composition of *Alphaproteobacteria*, *Gammaproteobacteria*, and taxa in the “Other” category (defined as any class with < 1% relative abundance across all time points), relative to all heterotrophic reads following the first phage infection (*t-test*, p < 0.05; Figure 9 a). Specifically, the relative abundance of *Gammaproteobacteria* decreased, while *Alphaproteobacteria* increased (Figure 9 a). This trend is also seen in experiments with an estuarine *Synechococcus* strain (Man et al 2024), suggesting that organic matter from phage lysis may favor *Alphaproteobacteria*. After the second attempted infection, resulting in no host lysis (Figure 6 b), we observed the opposite trend, with a significant increase in relative abundance of *Gammaproteobacteria* and a decrease in *Alphaproteobacteria* (*t-test*, p < 0.01; Figure 9 a). There was also a significant increase in relative abundance of *Betaproteobacteria* and heterotrophs in the “Others” category after the second infection (*t-test*, p < 0.01; Figure 9 a). This suggests that the heterotrophic community continues to change and reorganize at the class level, even in the absence of detectable host lysis.

**Figure 9.**
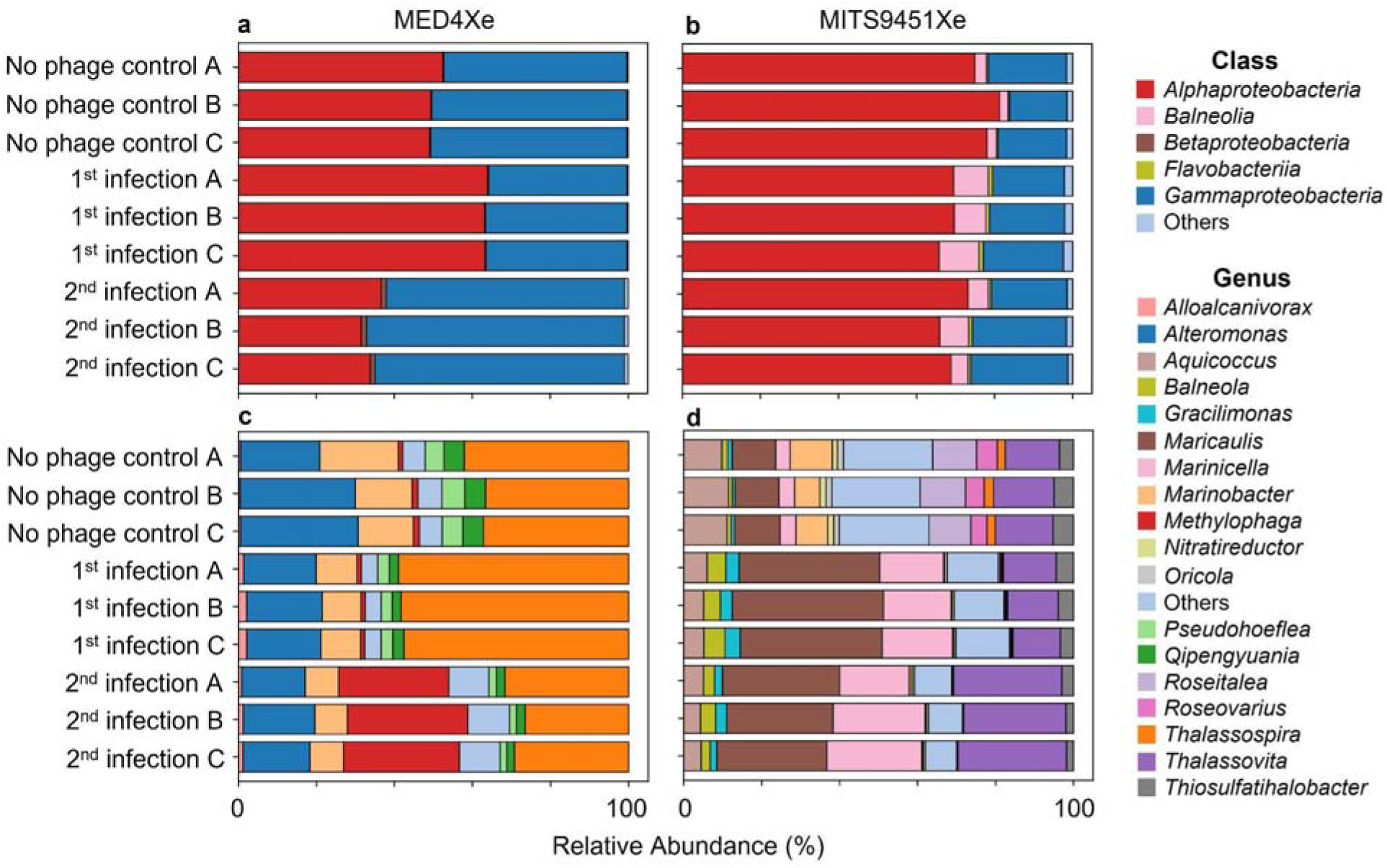
Heterotroph community composition changes following successive phage infections. Relative abundance of heterotrophic taxa as a proportion of total heterotrophic metagenomic reads were identified in xenic cultures before and after 2 rounds of phage infection. Data are classified to the class level for (a) *Prochlorococcus* MED4 and (b) *Synechococcus* MITS9451, and classified to the genus level for (c) *Prochlorococcus* MED4 and (d) *Synechococcus* MITS9451. Taxonomic classification was completed using ProSynTax (Coe et al. 2025) on shotgun metagenomic data generated by Illumina sequencing. Raw data is available in Supporting Information Table S2.

There were more nuanced and dynamic changes in the heterotrophic communities at the genus level. Significant shifts occurred in several low-abundance taxa, including those grouped into the “Others” category (defined as genera with < 1% relative abundance across all time points), as well as *Alloalcanivorax*, *Pseudohoeflea, Qipengyuania*, and *Thalassospira* (*t-test*, p < 0.05; Figure 9 c). Specifically, relative abundance of *Alloalcanivorax* and *Thalassospira* doubled or nearly doubled in relative abundance after the first infection, while all other classified genera declined (Figure 9 c). Notably, some strains of *Thalassospira* are capable of motility (López-López et al. 2002; Hütz et al. 2011), which may have conferred an advantage in reaching infection sites where chemoattractants were released, potentially explaining their marked increase in abundance. Following the second attempted infection, which resulted in no host lysis (Figure 6 b), relative abundance of *Marinobacter*, *Pseudohoeflea*, and *Thalassospira* significantly decreased, while *Methylophaga* and genera in the “Others” category significantly increased (*t-test*, p < 0.01; Figure 9 c). Thus, phage-induced lysis can drive rapid and pronounced restructuring of heterotrophic communities by releasing cellular contents that serve as new nutrient sources and by altering competitive dynamics. Even in the absence of lysis, changes still occur at both the class and the genus levels, reflecting the microbial taxa acclimating to changes in host physiology (e.g. growth rate and metabolism) and to the variations in the types and quantities of metabolites released by the host. Although *Alteromonas*, which was used in monoxenic *Prochlorococcus* infections in Figures 1, 2, 4, and 5, significantly increased in abundance during phage lysis in monoxenic cultures (Figure 3, Supporting Information Figure S1), in xenic cultures it decreased in relative abundance, suggesting that competition in a more complex community constrains its bloom despite its potential functional role in host recovery.

The heterotroph communities in *Synechococcus* MITS9451 cultures responded differently at the class level to those in the *Prochlorococcus* MED4 cultures following the first phage infection, with a significant decrease in *Alphaproteobacteria* and significant increase in *Balneola* and *Flavobacteria* (*t-test*, p < 0.05; Figure 9 b), two classes which were absent from xenic MED4 cultures. As with MED4, the second infection of MITS9451 culture did not result in host lysis; however, unlike MED4, it also did not lead to significant changes in community composition at the class level. At the genus level, half of the genera decreased in relative abundance, except for *Balneola*, *Gracilimonas*, *Maricaulis*, and *Marinicella* which increased (*t-test*, p < 0.05), and *Thalassovita* which remained unchanged (Figure 9 d). Following the second attempted infection, which resulted in no host lysis (Figure 6 c), *Maricaulis* declined in relative abundance, while *Thalassovita* increased (*t-test*, p < 0.05; Figure 9 d). Although the heterotrophic community composition in *Synechococcus* MITS9451 cultures differed from that of *Prochlorococcus* MED4, similar large shifts in abundance of multiple genera were observed after phage infection.

Thus, the structure of heterotrophic communities accompanying host cells, and by extension, the diversity of metabolites they produce, are altered by phage infection and continue to shift over sequential transfers of the community into fresh media. Notably, despite these ongoing changes, both cyanobacterial hosts used in our experiments were able to recover from the initial infection, implying that either community function was maintained in a way that continued to support host growth, or that changes had already occurred within the hosts themselves to prevent further infection.

## Conclusions

This work highlights the complexity of microbial and phage interactions in ocean ecosystems. Using axenic, monoxenic, and xenic *Prochlorococcus* MED4 as a simplified “model system”, we found that the presence of heterotrophs, whether a single species or community, facilitates *Prochlorococcus* recovery from phage infection (Figure 10). This interpretation applies across all *Prochlorococcus* phage:host systems tested, but not uniformly across *Synechococcus*:phage systems as detailed in Figure 10, highlighting important differences in host-specific responses.

**Figure 10.**
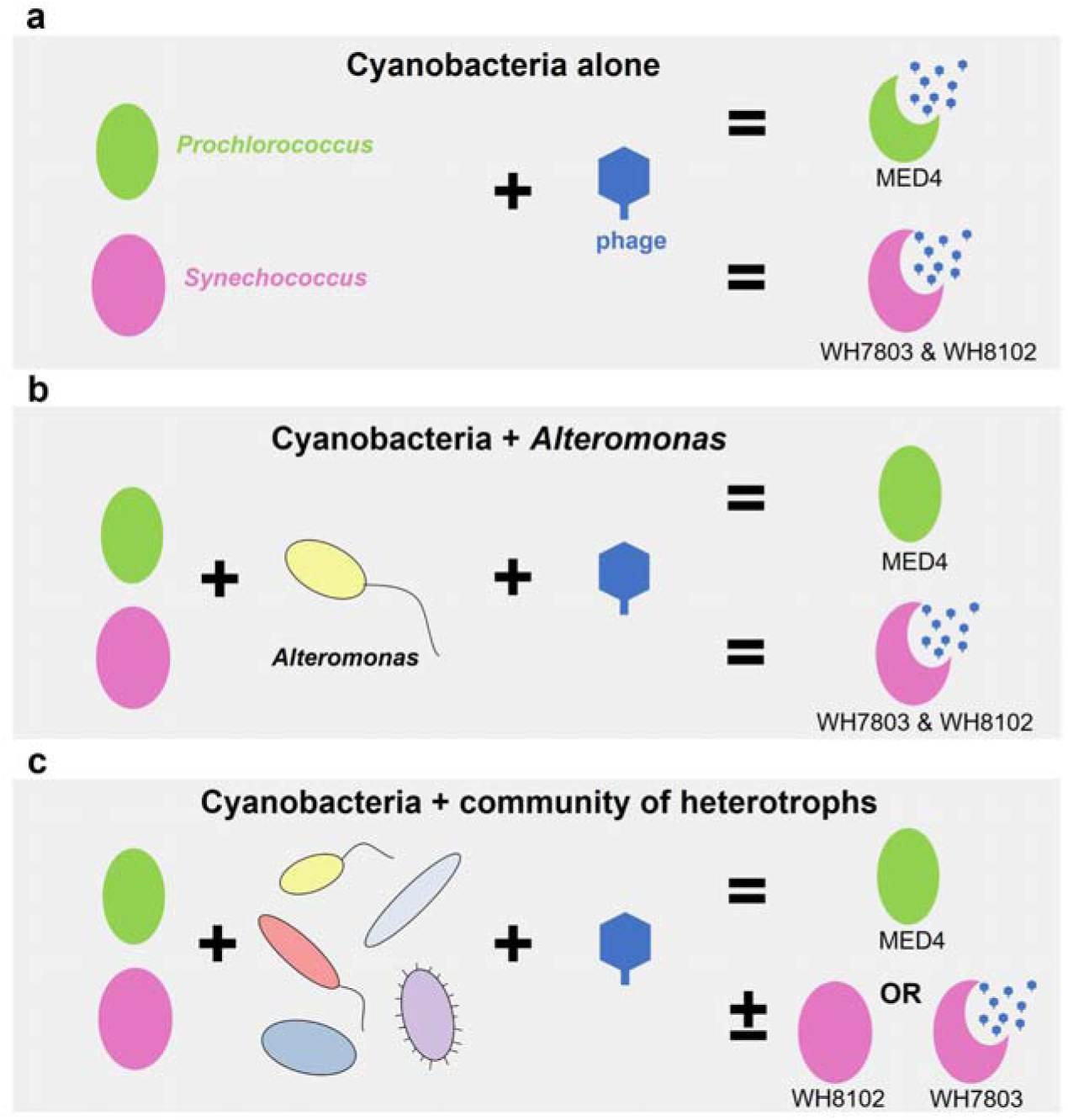
Phage:host outcomes depend on heterotroph presence and diversity. Cartoon schematic showing *Prochlorococcus* MED4 (green) and *Synechococcus* WH8102 and WH7803 (pink) cells (A) alone, (B) +*Alteromonas*, and (C) with a community of heterotrophs infected with phage (blue).

The broader message, however, is not the specific responses of individual phage:host:community combinations, but rather that biological context matters in interpreting biological systems. Traditional microbiology – and most of biology for that matter – often rely on single-species model systems, but these do not capture the interactions and cross-feeding that shape outcomes in more natural, complex communities. For example, responses differed when *Synechococcus* cultures were supplemented with one heterotroph versus multiple heterotrophs (Figure 5 and Supporting Information Figure S2), suggesting that specific metabolites can elicit distinct metabolic responses, including enhanced resistance to phages.

By incorporating ecological complexity, we gain a deeper understanding of the variability, resilience, and emergent properties of microbial systems. As properties of the system emerge over time they dictate the local environment which, in turn, applies selective forces on the system and determines its trajectory. The work presented here just scratches the surface, raising new questions about how host physiology (e.g., metabolic adaptability), community composition, and environmental context together determine recovery from phage infection.

## Acknowledgements

The authors thank both past and present members of the Chisholm Lab for support and comments. The authors also thank Trent LeMaster and Alexandrya A. Robinson for experimental assistance. This study was funded by the Simons Foundation (Life Sciences Project Award ID 337262, S.W.C.; SCOPE Award ID 329108, S.W.C.; Life Sciences Postdoctoral Fellowship Award ID 649394, S.M.K.). The authors declare no competing interests. The funding sources had no involvement in the study’s design, execution, or the decision to submit this work for publication.

## Data Availability

The metagenomes from this study were deposited in NCBI under Bioproject PRJNA1302459. All code used to process, analyze, and generate figures from the DNA sequencing data is available at: https://github.com/nhinvo/pro-het-phage-interaction.

## Supporting Information

**Supporting Information Table S1. Metagenomic samples before and after phage infection.** Samples from monoxenic and xenic *Prochlorococcus* MED4 and *Synechococcus* MITS9451 cultures collected prior to infection, after one round of phage infection, and after two rounds of phage infection, with corresponding sequencing accession numbers.

**Supporting Information Table S2. Sequence variants detected in xenic cyanobacterial cultures before and after phage infection.** Single nucleotide variants (SNVs), insertions, and deletions were identified in monoxenic and xenic *Prochlorococcus* MED4 and *Synechococcus* MITS9451 cultures prior to infection, after one round of phage infection, and after two rounds of phage infection. Variants are listed with the corresponding genome position and gene ID.

**Supporting Information Table S3. Relative abundance of heterotrophic taxa before and after phage infection.** Relative abundance of heterotrophic taxa, expressed as a proportion of total heterotrophic metagenomic reads, is shown for xenic *Prochlorococcus* MED4 and *Synechococcus* MITS9451 cultures before and after 2 rounds of phage infection. Data are presented at the community, class, and genus level based of taxonomic classification performed using ProSynTax ((Coe et al. 2025)) on Illumina shotgun metagenomic data.

**Supporting Information Figure S1.**
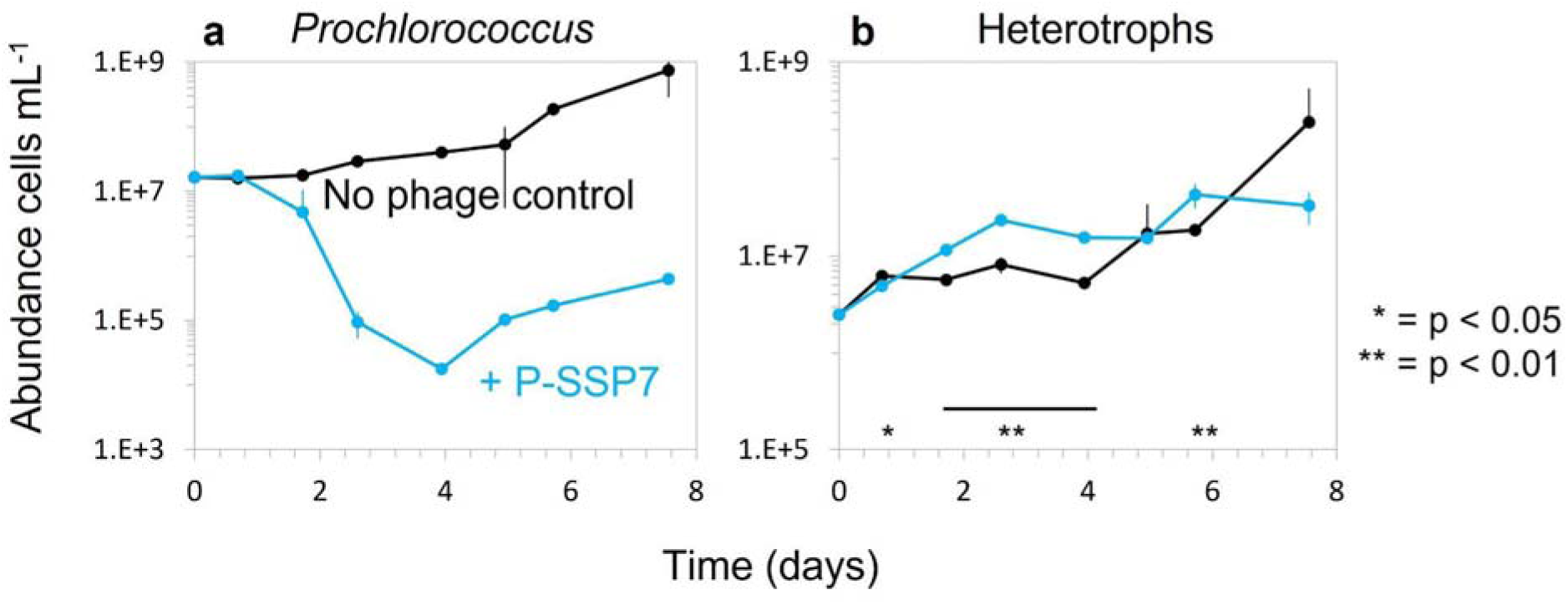
Heterotroph abundance increases during phage infection. Abundance of (a) xenic *Prochlorococcus* MED4 and (b) their associated community of heterotrophs with (blue) and without (black) podo-like phage P-SSP7 added at the onset of the experiment. Significance of heterotrophic growth differences (* = p <0.05, ** = p < 0.01) was determined by *t-test* and significance across multiple time points is indicated with a black line.

**Supporting Information Figure S2.**
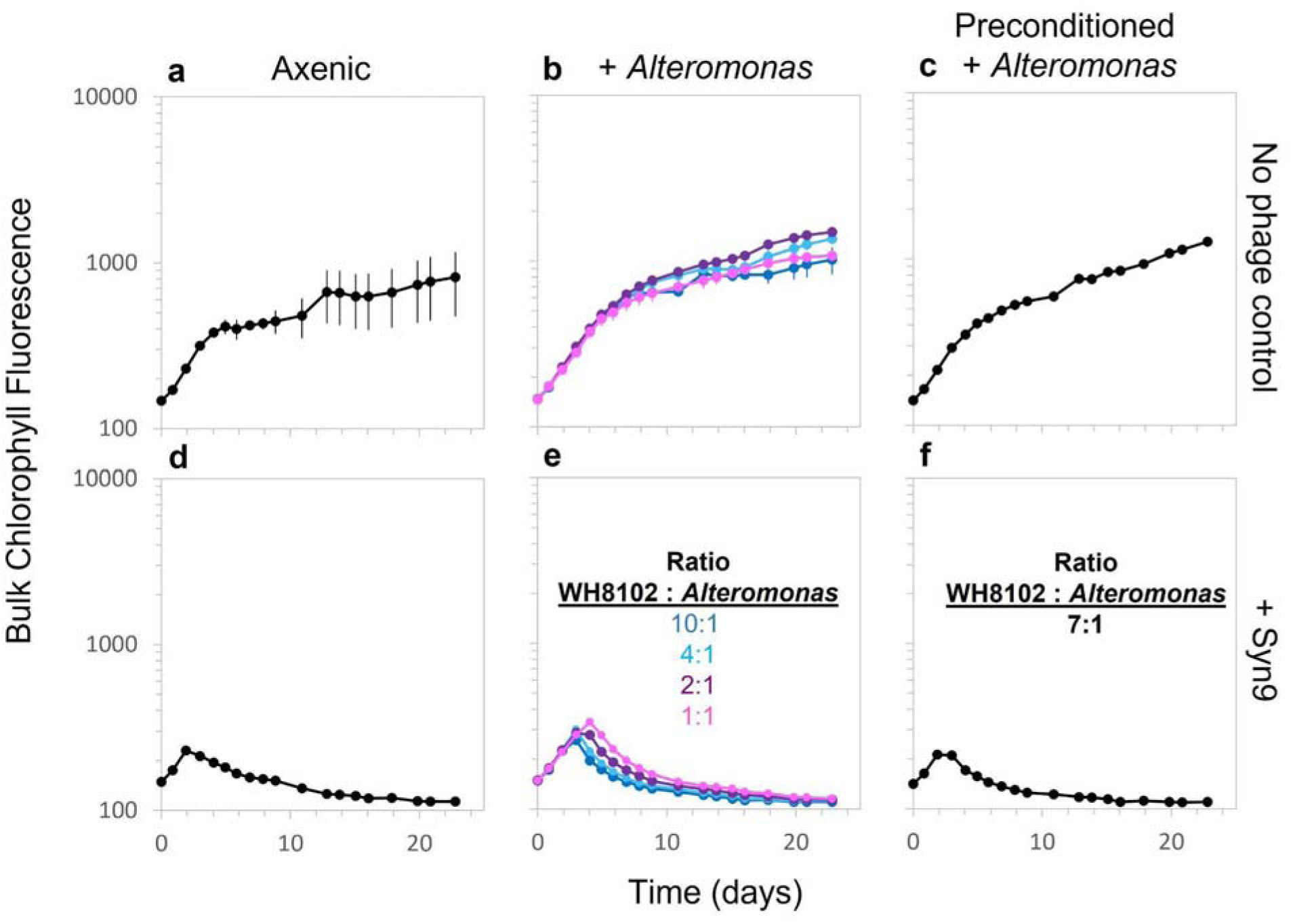
Presence of *Alteromonas* does not enhance *Synechococcus* recovery post-infection. Axenic and monoxenic (*+Alteromonas*) with varying initial ratios of *Synechococcus* to *Alteromonas*, and preconditioned monoxenic *Synechococcus* WH8102 cultures without (a,d) and with (b, c, e, f) cyanophage Syn9 added at the on-set of the experiment. Infection dynamics were monitored via chlorophyll fluorescence.

**Supporting Information Figure S3.**
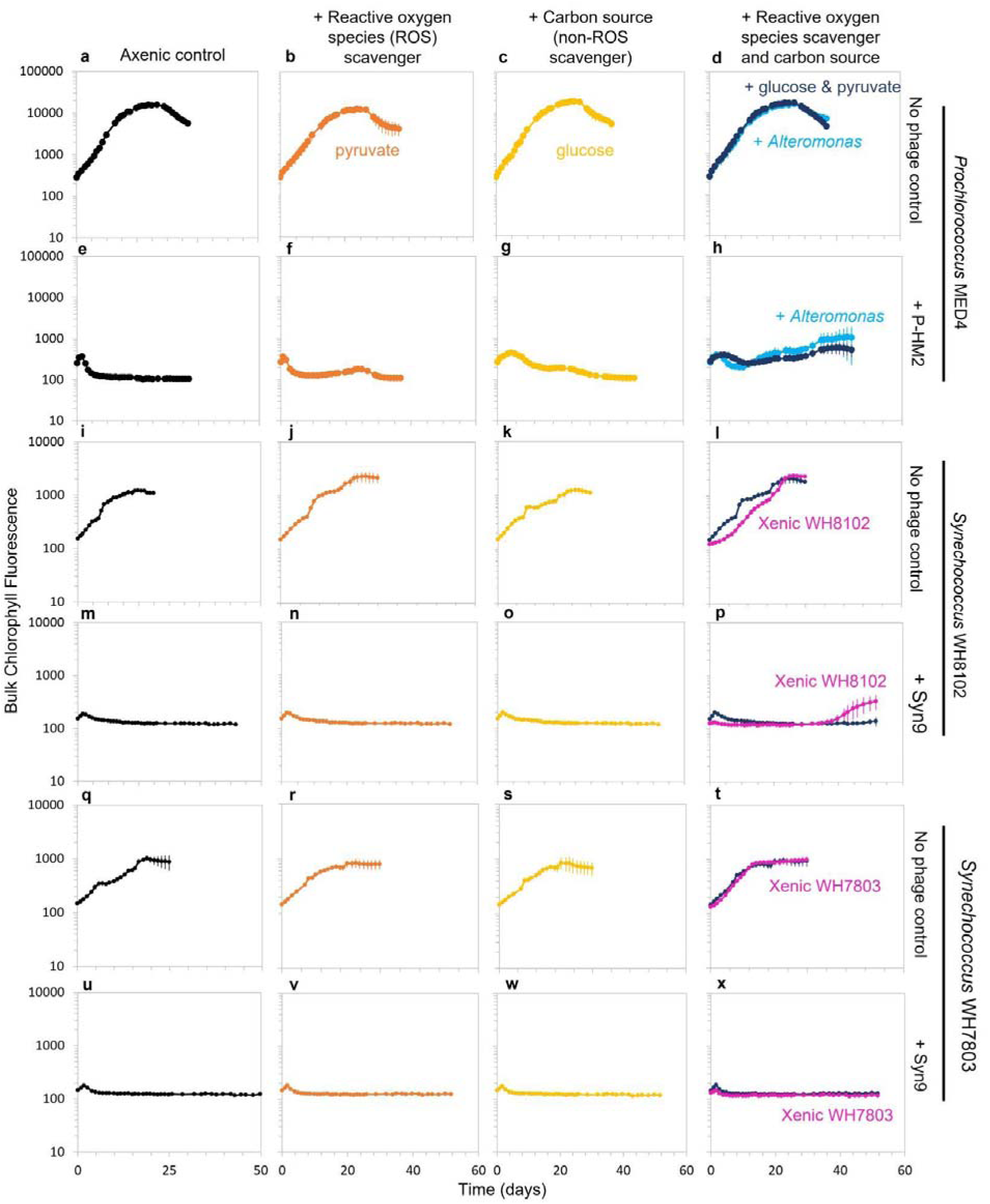
Reactive oxygen species (ROS) mitigation and carbon supplementation enable post-infection recovery in some phage:host combinations. Axenic cultures of *Prochlorococcus* MED4 and *Synechococcus* WH8102 and WH7803 were grown alone (a, e, i, q; black lines), with reactive oxygen species scavenger (ROS) pyruvate (b, f, j, n, r, v; orange lines), with organic carbon source glucose (c, g, k, o, s, w; yellow lines), or with a combination of glucose and pyruvate or heterotrophs (either +*Altermonas* or xenic, a community of heterotrophs) that can mitigate oxidative stress and provide carbon substrates (d, h, l, p, t, x; navy blue, aqua, or pink lines). The no phage control cultures were grown in parallel with cultures infected with either P-HM2 or Syn9 phages. All phage, substrates, or bacteria were added at the onset of the experiment, while heterotrophs in xenic cultures were already co-cultured with the host.

